# Tangential high-density electrode insertions allow to simultaneously measure neuronal activity across an extended region of the visual field in mouse superior colliculus

**DOI:** 10.1101/2021.06.12.448191

**Authors:** Jérémie Sibille, Carolin Gehr, Kai Lun Teh, Jens Kremkow

## Abstract

The superior colliculus (SC) is a midbrain structure that plays a central role in visual processing. Although we have learned a considerable amount about the function of single SC neurons, the way in which sensory information is represented and processed on the population level in awake behaving animals and across a large region of the retinotopic map is still largely unknown. Partially because the SC is anatomically located below the cortical sheet and the transverse sinus, it is technically difficult to measure neuronal activity from a large population of neurons in SC. To address this, we propose a tangential recording configuration using high-density electrode probes (Neuropixels) in mouse SC *in vivo* that permits a large number of recording sites (~200) accessibility inside the SC circuitry. This approach thereby provides a unique opportunity to measure the activity of SC neuronal populations composing up to ~2 mm of SC tissue and characterized by receptive fields covering an extended region in the visual field. Here we describe how to perform tangential recordings along the anterior-posterior and the medio-lateral axis of the mouse SC *in vivo* and how to combine this approach with optogenetic tools for cell-type identification on the population level.

## Introduction

The superior colliculus (SC) is a multisensory structure that receives direct visual sensory inputs from retinal ganglion cells and that plays multiple important roles in visually guided behaviors. While the SC has been studied extensively on the single neuron level (Ahmadlou and Heimel, 2015; Ahmadlou et al., 2017; Gale and Murphy, 2018; 2016; 2014; Inayat et al., 2015; Lee et al., 2020; Shi et al., 2017; L. Wang et al., 2010), the role of its neural circuits and specific cell-types under *in vivo* conditions, in awake behaving animals, remains elusive. Recently, its role in sensory processing and also its participation in higher cognitive functions regained attention because modern genetic tools and sophisticated behavioral measurements became available (Basso and May, 2017; Zhao et al., 2014; Basso et al., 2021; Cang et al., 2018; De Franceschi and Solomon, 2018; Evans et al., 2018; Ito and Feldheim, 2018; Reinhard et al., 2019; Sans-Dublanc et al., 2021; Savier et al., 2019; Villalobos et al., 2018; Zhang et al., 2019). However, it is still not well understood how sensory information from the complex and dynamic visual environment of the entire visual field (Qiu, et al., 2021) is represented at the population level within SC and what role the different cell-types play in extracting and representing behaviorally relevant information (Gale and Murphy, 2014; Hoy et al., 2019; Sans-Dublanc et al., 2021).

One reason for our limited understanding about the neural computations on the population level is the anatomical location of the SC below the cortex and the transverse sinus. Therefore, recording simultaneously from large populations of SC neurons remains technically challenging. Two approaches have been developed that allow recording from multiple SC neurons simultaneously in mice *in vivo*. Multi-electrode silicone probes are regularly employed to record from SC neurons by inserting the probe into the SC tissue by way of the cortical sheet (Ahmadlou et al., 2017; Ahmadlou and Heimel, 2015; Lee et al., 2020). The benefit of this approach is that the cortex does not need to be removed to provide access to the SC. Consequently, the remaining cortex enables the study of the SC in awake behaving animals. In addition, multi-electrode arrays record neuronal activity simultaneously across the different SC layers, thereby providing layer specific information on the location of the probe and neurons within the SC via current source density analysis (Ahmadlou et al., 2017; Zhao et al., 2014). While this vertical recording approach has provided important insights into the processing of visual information on the single neuron level (Ahmadlou et al., 2017; Ahmadlou and Heimel, 2015; Ito et al., 2017; Lee et al., 2020), it has limitations in that the activity of the population of SC neurons which can be simultaneously recorded has receptive fields located in an overlapped and limited location in the visual field. Here it is important to note that visually guided behaviors depend on the simultaneous integration of information across the entire visual field (Hoy et al., 2019; 2016; Yilmaz and Meister, 2013), therefore, the study of such behaviors requires monitoring the collective activity from a population of neurons within the horizontal plane of the SC layers to reveal the underlying circuit mechanisms.

The other approach to study the activity of neural populations across the horizontal dimension of neural circuits is two-photon calcium imaging. While this technique is the method of choice to record from large populations of cortical neurons across the dorsal surface (de Vries et al., 2020; Froudarakis et al., 2014; Kim et al., 2016; Stringer et al., 2019) and simultaneously across a large extent of the retinotopic map in the visual cortex (Zhuang et al., 2017), its applicability to measure neuronal activity across the SC surface is limited due to its subcortical location. Recently, several studies have developed methods that overcome this challenge (Barchini et al., 2018; Feinberg and Meister, 2015; Inayat et al., 2015; Li et al., 2020). Feinberg et al. have developed a silicone plug with which the transverse sinus can be pushed anteriorly to expose a small area of the posterior SC (Feinberg and Meister, 2015). The silicone plug is transparent and allows to study the activity of SC neurons by two-photon calcium imaging. This technique has revealed important new insights into SC processing and the circuits functional organization (Feinberg and Meister, 2015; Li et al., 2020). However, this approach has some limitations as it allows imaging from only a small part of the posterior SC. Studying the anterior part of the SC, that represents the frontal and upper visual field, requires to remove the overlying cortex to expose the SC surface (Barchini et al., 2018; de Malmazet et al., 2018; Inayat et al., 2015; Savier et al., 2019) or to employ transgenic mouse lines that lack a developed cortex (Gribizis et al., 2019). However, both procedures require removing an essential part of the mammalian brain, the cortex, and therefore prevent the study of population activity in SC under natural conditions. Taken together, we are still limited in methodologies that would allow the measurement of large neural populations across the entire retinotopic map of SC *in vivo*.

Here we describe a novel approach for recording neuronal activity across a large extent of the SC circuitry *in vivo*. This method enables recording large populations of neurons with their receptive fields (RFs) covering an extended region of the visual space. The key for achieving this large-scale measurement is the combination of high-density electrode arrays (Neuropixels probes (Jun et al., 2017)) together with an insertion angle tangentially to the SC surface. High-density electrodes allow to sample neuronal activity at an unprecedentedly large scale, across several millimeters, and high spatial resolution. The tangential probe insertion makes it possible to study neuronal activity across the horizontal dimension of neural circuits (Kremkow et al., 2016), thereby providing a high yield of simultaneously recorded neurons from within the same brain structure. Here we demonstrate that combining high-density probes with tangential insertions is a promising technique for large-scale, *in vivo* recordings from SC neuronal populations, across a substantial extent of the visual field. The aim of this article is to provide an outline on how to perform these tangential recordings in mouse SC. Specifically, we will provide a guide on how to perform these insertions along the anterior-posterior (AP) axis and the medio-lateral (ML) axis of SC and how to adjust and optimize probe placement during the experiment using functional measurements. Both recording configurations are similar in their implementation and complementary as they sample distinct axes of the visual field. We also show that these recording configurations are effective ways to characterize the functional organization within SC. At last, we will illustrate how to combine the tangential insertions with precise optic fiber placement for optogenetic identification of populations of SC cell-types.

## Material and Methods

In the following, we will describe how to target a large population of SC neurons via tangential Neuropixels probe implantations.

### Surgery

All experiments were pursued in agreement with the local authorities upon defined procedures (LAGeSo Berlin - G 0142/18). During all experiments, maximum care was taken to minimize the number of animals. Adult male mice (C57BL/6J) of 2-7 months old were used from the local breeding facility (Forschungseinrichtung für Experimentelle Medizin, Charité Berlin) and Charles-River, Germany. For optotagging experiments, VGAT-ChR2 mice of 5-7 months of age were used.

For anesthesia induction, animals were placed in an induction chamber and exposed to isoflurane (2.5%, CP-Pharma) in oxygen (0.8 L/minute). Once anesthetized, the surgery was performed in a stereotactic frame (Narishige) with a closed-loop temperature controller (FHC-DC) for monitoring the animal’s body temperature. The isoflurane level was gradually lowered during surgery (0.7-1.5%) while ensuring a complete absence of vibrissa twitching or responses to tactile stimulation. During surgery, the eyes were protected with eye ointment (Vidisic). Mice were head-fixed in the frame, and the skull was exposed. The head was aligned along the anterior-posterior axis and marks were made for the craniotomy at 600-1500 μm ML and 0-1500 μm AP from Lambda. For optotagging experiments, an additional mark for the optic fiber placement was made at 3500 μm AP, 700-900 μm ML to Bregma. A headpost was placed on the skull and implanted with dental cement (Paladur). During this step, a silver wire (AG-10W, Science Products) was attached within the dental cement based recording chamber which was surrounding the craniotomy. Here special attention was taken to keep the recording chamber on the on the posterior or lateral side low enough to make room for the shallow angle probe implantation. When dry, craniotomies were made at the marked positions with a dental drill (Johnson-Promident). To allow for the later probe insertion at a horizontal angle between 15° and 25°, a small part of the skull at the posterior part of the craniotomy was slightly thinned using the drill to ensure a smooth insertion.

### Electrophysiological recordings and visual stimuli

Electrophysiological SC recordings were performed using Neuropixels probes (Jun et al., 2017). Data was acquired at 30 kHz using the Open Ephys acquisition system (Siegle et al., 2017). Multi-unit activity (MUA) analysis (see below) was carried out using customized Python scripts.

Visual stimuli were generated in Python using the PsychoPy toolbox (Peirce, 2008) and presented on a spherical dome screen that allows to present visual stimuli across a large part of the visual field (Denman et al., 2018; 2017) (Fig. 1a/b). The visual stimuli were displayed on a calibrated and warped projector image (EPSON projector, refresh rate = 60 Hz, mean luminance = 110 cd/m^2^). The projector image was reflected into a plastic spherical dome screen (EBrilliantAG, IP44, diameter = 600 mm) upon a spherical mirror (Modulor, 0260248). The inside of the plastic dome was covered with a layer of broad-spectrum reflecting paint (Twilight-labs). Calibration of the dome warping was done with the Meshmapper software (Paul Bourke). In a subset of experiments, we used a calibrated LCD screen (Dell, refresh rate = 120 Hz, mean luminance = 120 cd/m2).

**Figure 1:**
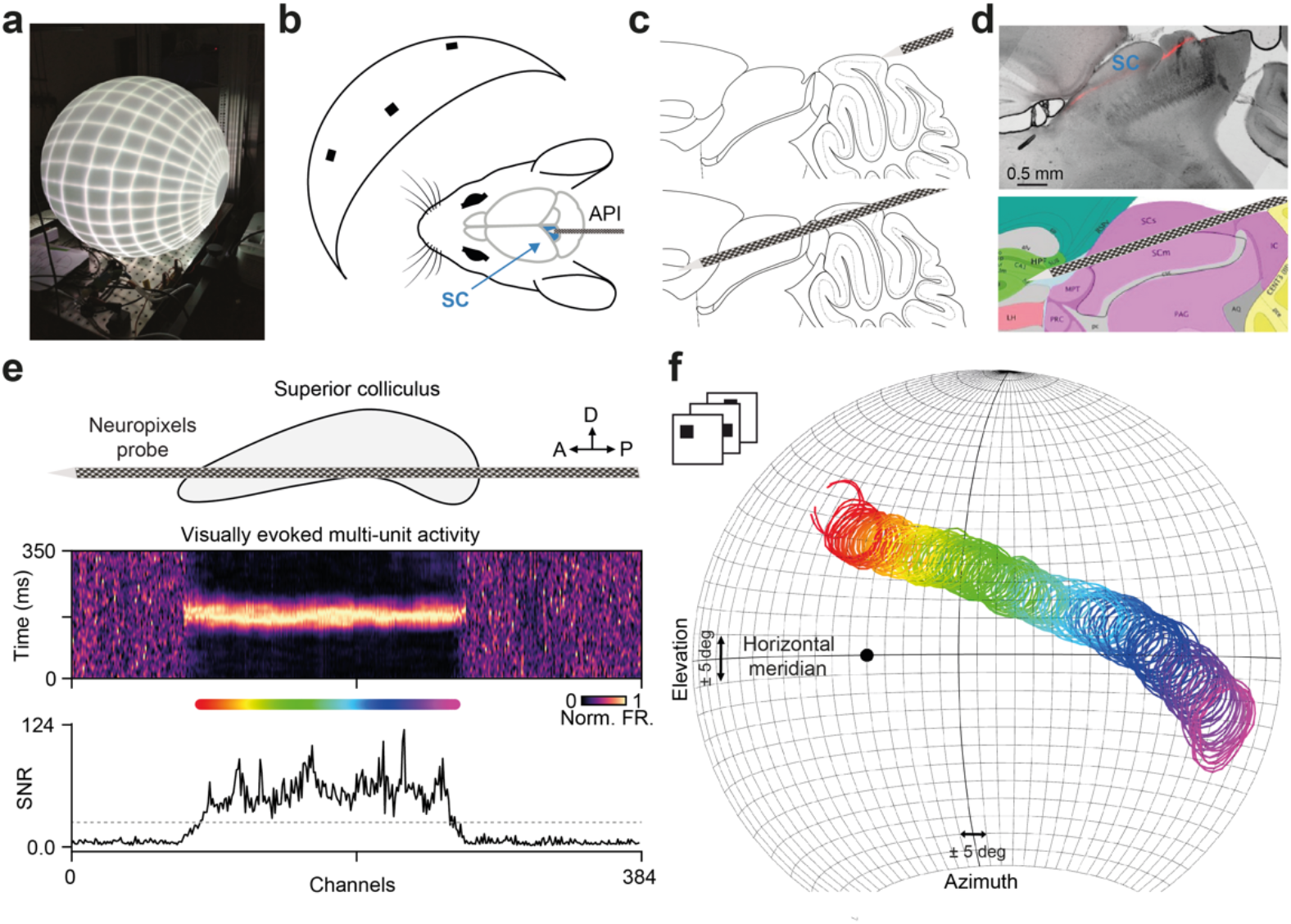
Tangential recordings in the mouse superior colliculus (SC) along the anterior-posterior axis. (**a**) Spherical dome visual stimulation system. (**b**) Schematic of the animal position in the dome together with the tangential anterior-posterior insertion (API) targeting the upper visual layers of the SC. (**c**) Schematic of the initial and final probe position. (**d**) Sagittal section containing the DiI staining (red) of a Neuropixels electrode track (top) and the corresponding location in the Allen Institute Common Coordinate Framework (CCF, bottom). (**e**) Visually evoked multi-unit activity (MUA) to a sparse noise stimulus is used to confirm the probe placement within the SC during the experiment. Recording sites located inside SC respond strongly to the visual stimulus. Shown are the peri-stimulus time histograms (PSTHs, middle) and the signal-to-noise ratio along the channels (SNR, bottom). Note that ~200 recording sites are located within the SC, indicated by the rainbow colorbar. (**f**) Receptive fields (RFs) from recording sites with high SNR values (threshold shown as gray line in e) projected onto the spherical coordinates of the dome. The colors of the RFs correspond to the location within SC shown in e. Note the large extent of retinotopy captured by the API. The black dot shows the position of the mouse nose.

### Confirmation of probe placement by receptive field analysis

A real-time analysis of the RFs is crucial for the confirmation of the probe in the area of interest within the SC and the corresponding visual field. Currently, it is still not possible to perform online spike detection during the recording in the OpenEphys acquisition system due to the high channel count of Neuropixels probes. However, to have an estimate of the recorded retinotopy covered by the implantation during the experiment, we implemented a semi-online RF analysis by presenting a short sparse noise stimulus (stimulus duration ~5 min) and directly afterwards performing a RF estimation on the multi-unit activity (analysis duration ~5 min).

To measure RFs, we presented locally sparse noise (LSN) stimuli (light squares on dark background) with target squares 5-15° in size and updated 10 Hz (50-100 ms per LSN frame). To minimize the spatial correlation, the target squares in an LSN frame were designed to be separated by at least seven LSN squares.

The RFs were estimated from the MUA using spike-triggered average (STA). To obtain the MUA, we first applied the common average referencing to the raw signals. Then, we bandpass filtered the processed signals using a Butterworth filter (order of 2, 0.3 to 3 kHz) and detected the spiking events that have amplitudes larger than four times the standard deviation of the filtered signals. Due to the high channel count of the Neuropixels probe, these steps are computationally expensive. In order to perform these analysis steps within a few minutes, we used the Python parallel processing package “Joblib” (https://joblib.readthedocs.io/en/latest/) to extract the MUA across multiple channels in parallel. Having access to the RFs of all visually driven channels during the recording allowed us to assess the recording location within the SC circuitry by means of RF locations in the visual field (Fig. 2). The Python code for performing the RF analysis is available from the corresponding author on request.

**Figure 2:**
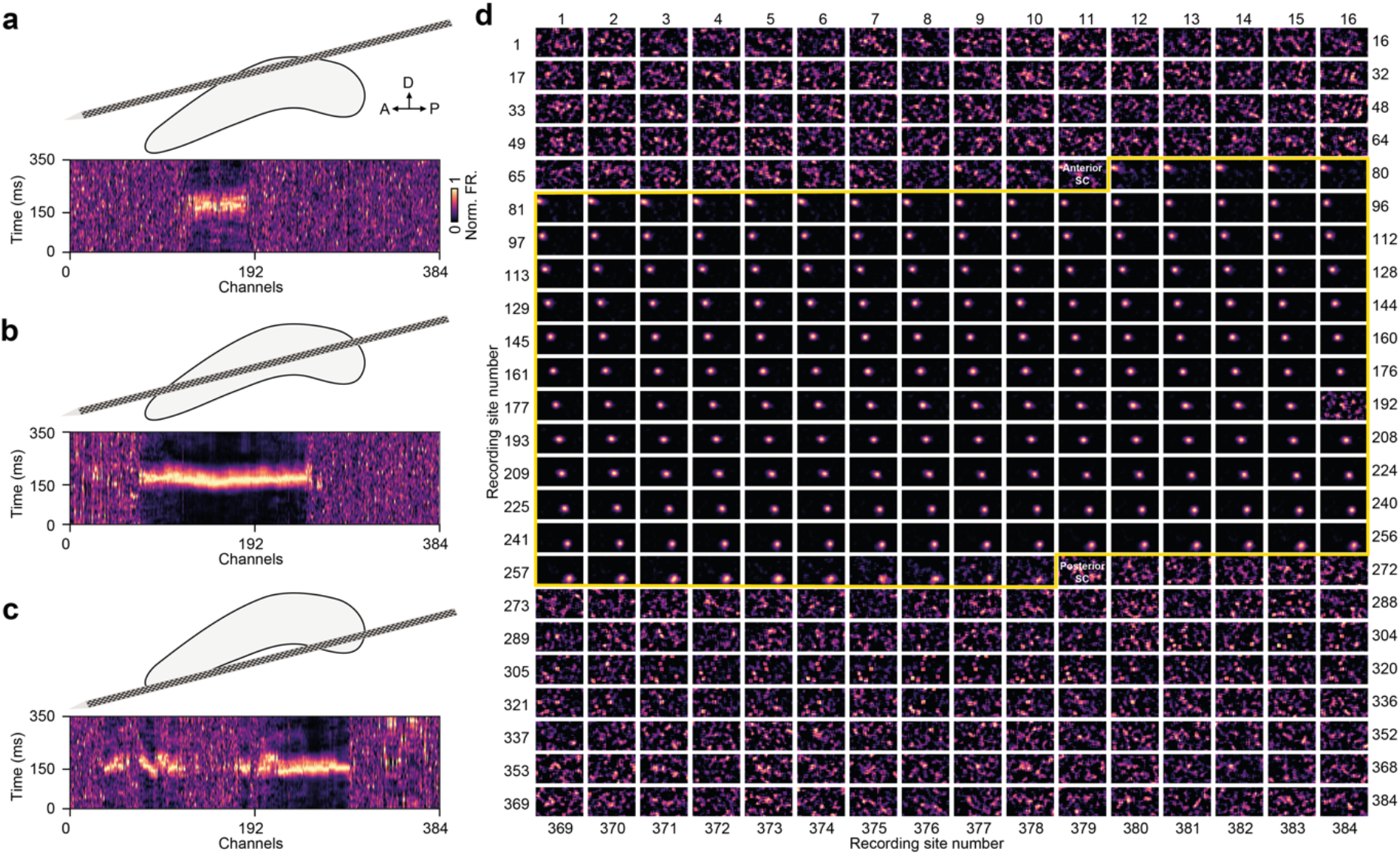
Functional confirmation of probe placements in the upper visual layers of the mouse SC. (**a-c**) Schematic of the putative probe placement (top) and its corresponding MUA PSTH (bottom). (**a**) Visually evoked activity in only a few continuous recording sites indicates an excessively superficial insertion. Relocation of the probe more ventrally could cover a larger visually driven area. (**b**) Visually evoked activity across a large number of consecutive recording sites indicates a correct placement of the tangential insertion in the upper visual layers of the SC. (**c**) Visually driven activity in two discontinuous regions on the probe suggests an excessively ventral location of the probe. Relocating the probe towards more dorsal could increase the area of visually evoked activity. (**d**) Representative example of receptive fields (RFs) estimated by spike-triggered averaging of MUA during the experiment. Note the large number of visually driven channels and the continuous changes of retinotopic positions of their RFs.

### Histology

The reconstruction of the electrode track is required for post-hoc verification of the exact recording site location inside the brain. To that end, the probe was removed, coated with a fluorescent dye (DiI, Abcam-ab145311) and re-inserted in the same location. Subsequently, the animal was terminated with an excess of isoflurane (>4%) or a subcutaneous injection of a Ketamine/Xylazine (Ketamidor 1 g/mL, Rompun 2%) and perfused in PBS and 4% paraformaldehyde (PFA). Brains were post-fixed overnight in 4% PFA, sliced (100μm) on a vibratome (Leica VT1000S), and sections were mounted using a mounting medium containing DAPI. Slices were imaged on a fluorescence microscope using a 2.5x objective with their respective excitation length to visualize DiI and DAPI. Subsequently, the brain slices were aligned to the Allen Institute Common Coordinate Framework (CCF) (Q. Wang et al., 2020), and the location of the electrode track was identified (Fig. 1d).

### Tangential recordings in mouse superior colliculus along the anterior-posterior axis

To study the processing and representation of sensory information in neural circuits within neural populations, a high number of simultaneously recorded neurons is required. With this aim, we set out to record from a population of neurons in the visual layers of the SC using a tangential recording configuration along the anterior-posterior axis. The superficial part of the SC along this axis is relatively flat for about 1.5-2 mm (Huberman et al., 2008) (Fig. 1d), and curved only towards its posterior end. With properly targeting the flat part of SC, this recording configuration allows to capture neuronal activity from a large part of the SC circuitry using the high and dense sampling of the recording sites on the Neuropixels probe, both in anesthetized and awake mice.

In order to properly align the probe parallel to the SC surface, an insertion angle between 15° and 25° relative to the horizontal plane is required. The AP insertion enters the brain above the cerebellum >1 mm posterior to Lambda in order to pass right below the transverse sinus (Fig. 1c, top). Extra care has to be taken to avoid damaging the sinus during probe insertion. Consequently, it is advised to extend both the craniotomy and the recording chamber, as posterior as possible, to allow the probe shank to pass below the transverse sinus for targeting the superficial SC layers along the AP axis.

The shallow insertion angle is prone to slightly bending the shank of the Neuropixels probe while the probe enters the brain. Therefore, monitoring the position and shape of the Neuropixels probe is a crucial factor for successfully targeting the superficial SC layers and avoiding breaking the probe shank. Furthermore, the surface tension of the grounding solution can change the insertion angle, by pulling the tip towards the surface of the solution. To avoid that, the grounding solution should be removed during the insertion procedure, in particular during the first few mm of insertion. After this initial insertion step, the probe is slowly inserted to a depth of >4 mm (Fig. 1c, bottom). Upon reaching the target depth the probe is withdrawn for about 20 to 50 μm in order to release accumulated mechanical pressure. Furthermore, the probe was allowed to settle for ~10-20 minutes before assessing and confirming its location within the upper visual layers of the SC.

An important step for achieving proper probe placement and large population recordings in SC is to validate the probe position during the experiment by functional measures, i.e. visually evoked activity. Due to the retinotopic organization of the SC (Mrsic-Flogel et al., 2005; Cang et al., 2008), the RF measure can be used to estimate the location within the SC tissue (Fig. 1f, Fig. 2a and see “Confirmation of probe placement by receptive field analysis”). Furthermore, the number of continuous channels showing visually driven activity provides another important information on tissue location. Visually driven activity on only a few channels reveals non-optimal placement that could indicate a more superficially located probe covering only a small dorsal part of SC (Fig. 2a, top) or a too steep insertion angle. Two visually driven regions located at the anterior and posterior parts of the probe and separated by a region of non-visually driven channels, indicate a more ventral recording position covering two distinct parts of the SC (Fig. 2a, bottom). In both cases, the probe could be relocated to increase the number of recording sites within the SC tissue.

An optimal located AP insertion, that targets the flat part of SC (Fig. 1d, Fig. 2a, middle, Fig. 2b), can cover up to almost 200 recording sites with visually driven activity (Fig. 1e, Fig. 2b), corresponding to almost 2 mm within the SC. Due to this large-scale sampling of the SC tissue, the corresponding RFs extend over a large region of the visual field (Fig. 1f, Fig. 2d). Since the nasal-temporal axis of the visual field is mapped onto the anterior-posterior axis in SC, and the dorsal-ventral visual field axis onto the medio-lateral axis in SC (Dräger and Hubel, 1976), the RFs in the AP insertion predominately change position along the azimuthal plane (Fig. 1f). The visual field that can be covered with this method can range up to 140° in azimuth and 40° in elevation (Fig. 1f).

Regular LCD screens, located close to the eye of the mouse, are commonly used to present visual stimuli across a large area of the visual field (e.g, de Vries et al., 2020; Zhuang et al., 2017). However, LCD screens are not curved; therefore, they introduce spatial distortions for stimuli that are presented outside the center of the screen. A more precise approach for presenting visual stimuli simultaneously across the large visual field covered by our recording technique is to present the stimuli in a visual dome (Denman et al., 2018; 2017) (Fig. 1a). The surface of the dome, onto which the stimuli are projected, is curved; thus, presenting visual stimuli without major spatial distortions across the whole extent of the visual field of mice is possible.

In summary, the tangential anterior-posterior insertion of high-density electrodes into the SC introduced here allows to record neuronal activity from a large extent of the SC circuitry with RFs covering a substantial part of the visual field along the nasal-temporal axis.

### Tangential recordings in mouse superior colliculus along the medio-lateral axis

While the API allows to cover a wide range along the azimuthal plane, it only provides limited access to the elevation axis. Having access to the elevation axis would be of great interest as it has been shown that visual stimuli are processed differently at distinct retinotopic locations across the upper and lower visual fields in the retina (Baden et al., 2020; Sabbah et al., 2017; Szatko et al., 2020; Warwick et al., 2018). However, less is known about how the upper and lower visual fields are represented in the SC (de Malmazet et al., 2018). Since the elevation axis of the visual field changes along the medio-lateral SC axis, we established a medio-lateral insertion (MLI) with the overall insertion procedure being similar to the API described above. The main difference is that for MLI, the Neuropixels probe is inserted tangentially through the contralateral cortex into the SC (Fig. 3a/b).

**Figure 3:**
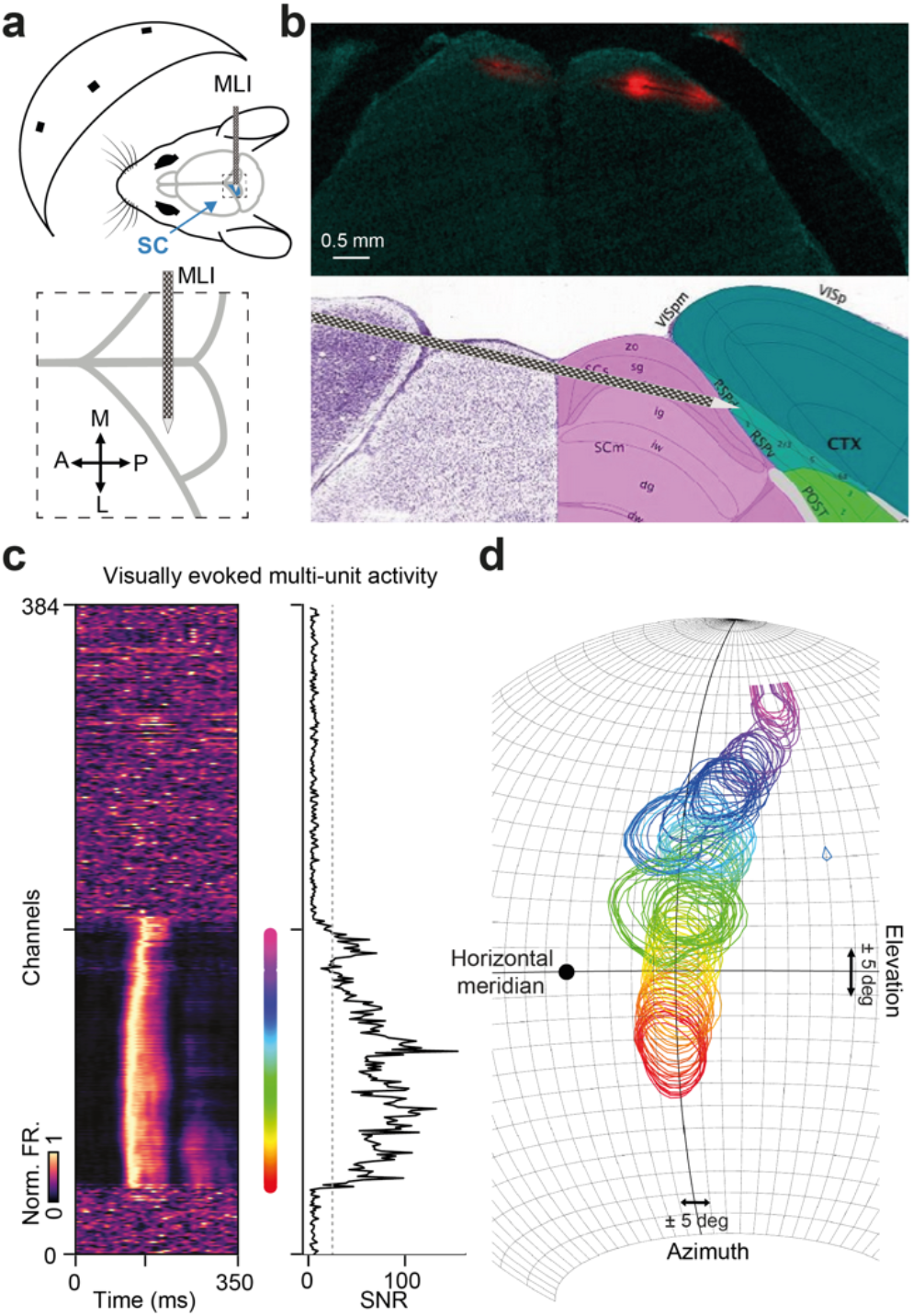
Tangential recordings along the medio-lateral (ML) axis of the SC. (**a**) Schematic of probe position for the ML configuration (top) with a zoom on the tangential ML insertion (MLI, bottom). (**b**) Coronal section of the MLI probe track (top) and the corresponding location in the Allen CCF (bottom). (**c**) Visually evoked multi-unit activity functionally confirms the electrode placement inside the upper visual SC layers. Same format as Fig. 1e. (**d**) Receptive fields (RFs) in the MLI. Note that RF locations in the MLI cover a large extent along the elevation axis, including positions in the upper visual field.

The MLIs are performed using a 20° to 30° angle relative to the horizontal plane. As for the API, the recording chamber should be sufficiently large such that the Neuropixels probe can be positioned and inserted at the desired angle. One challenging aspect of this recording configuration is the long distance between the first contact of the probe with the cortical tissue and the actual location of the SC in the contralateral hemisphere. Therefore, functional confirmation of probe placement using RF mapping is required and strongly advised (see above and Fig. 2a).

An optimal located MLI (Fig. 3c) can cover up to 200 visually driven recording sites (Fig. 3c) which corresponds to almost 2 mm within the SC along the medio-lateral axis. Because of the dorsal-ventral visual field axis onto the medio-lateral axis in SC, the RFs in the MLI predominately change position along the elevation plane (Fig. 3d). The visual field that can be covered and recorded with this method can range up to 40° in azimuth and 100° in elevation (Fig. 3d). Since RFs are even located in the upper part of the upper visual field, stimulus presentation using a visual dome is recommended.

Taken together, the tangential medio-lateral insertion allows to record neuronal activity from an extended region in the SC with RFs covering a large part of the visual field along the dorsal-ventral direction, including positions in the upper part of the upper visual field.

### Functional organization of the mouse superior colliculus

The tangential recordings are one method of choice for studying the functional organization along the horizontal dimension of neural circuits (Hubel and Wiesel, 1977; Kremkow et al., 2016). It has been reported that SC is organized in maps of visual space, orientation, and motion direction (Ahmadlou and Heimel, 2015; Feinberg and Meister, 2015; Li et al., 2020), but see (Chen et al., 2021). To demonstrate that tangential high-density recordings in mouse SC are elegant techniques for characterizing its functional organization, we studied how orientation and direction tuning changes along the recording track. To that end, we presented moving bars in 12 motion directions to characterize the tuning properties for each recording site from the multi-unity activity (Fig. 4). Orientation and direction preference were extracted from fitting a von Mieses function (Kremkow et al., 2016) to the tuning curve which was estimated from trial averaged responses (Fig. 4a, n = 30 trials).

**Figure 4:**
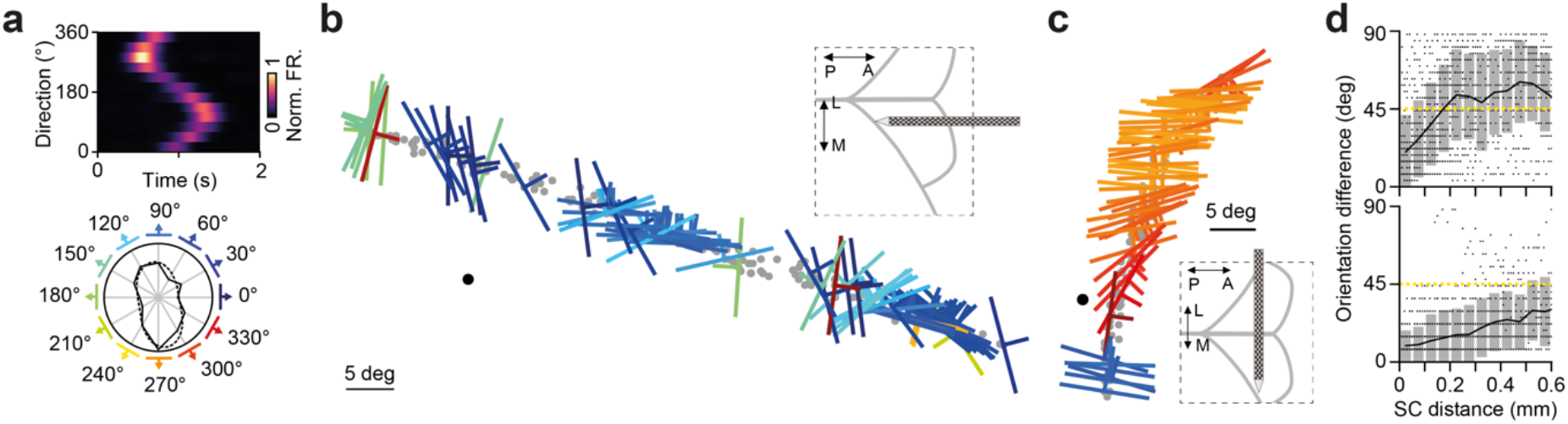
Topographic organization of orientation and direction in mouse SC revealed by tangential recordings. (**a**) Orientation and direction tuning were measured using moving bars. The evoked responses to moving bars of various directions (top) are used to calculate the tuning curve shown in the polar plot (bottom). (**b**) Illustration of the functional organization along an API (inset top right). Shown are receptive field locations (gray dots) of recording sites with high signal-to-noise values. The preferred direction and orientation are shown by the color and the angle of the bars. The colors of the bars correspond to a. (**c**) Functional organization along an MLI. Note the systematic change in orientation and direction preference along the elevation axis. Same format as in b. (**d**) Orientation difference as a function of channel distance for the example shown in b (top) and c (bottom). Note that nearby locations are similarly tuned. The yellow dashed line represents random uniform tuning.

Our data confirmed that orientation and direction preference are systematically organized along the horizontal dimension of SC, at least on the multi-unit level (Fig. 4b-c). Both along the anterior-posterior/azimuth axis (Fig. 4b, left; Fig. 4d, top) as well as along the medio-lateral/elevation axis (Fig. 4b, right; Fig. 4d, bottom). This exemplifies that the tangential high-density electrode approach is suitable for characterizing the functional organization of the SC.

### Optogenetic identification of large populations of SC cell-types

The tangential implantations provide an opportunity to record from a large neural population within the SC. As a next step, we tested whether those tangential insertions can be combined with optogenetic tools to identify large populations of defined cell-types in SC. To that end, we established a configuration where the optic fiber is placed orthogonal to the Neuropixels probe to optogenetically-activate a wide range of inhibitory cells in SC (Fig. 5a).

**Figure 5:**
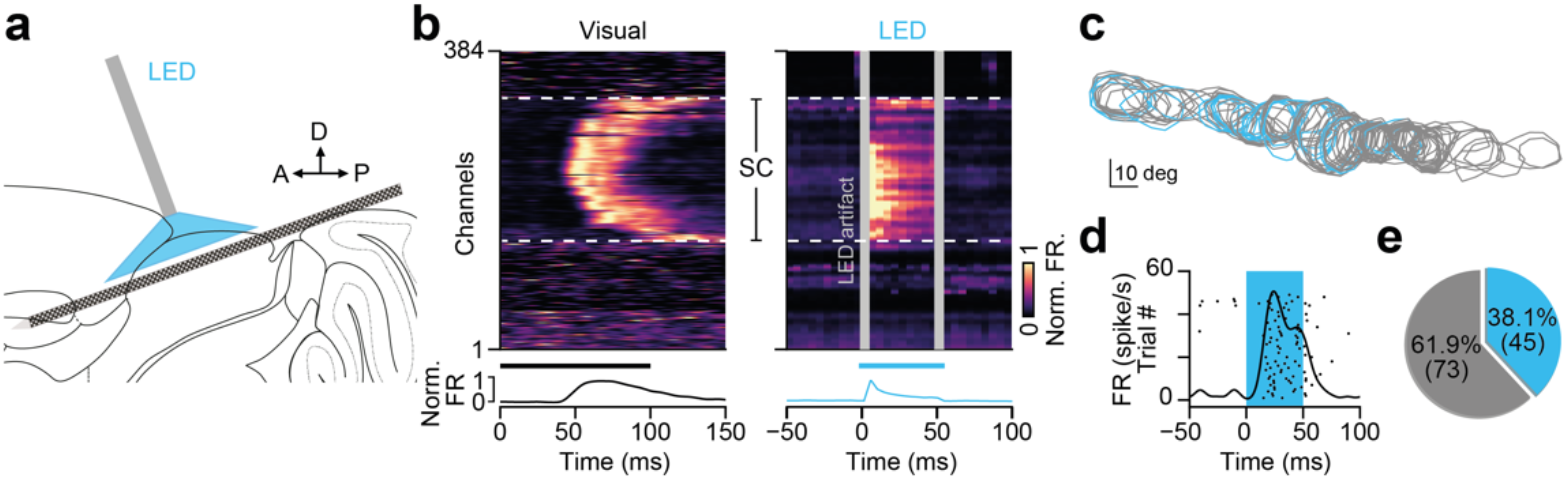
Combining optogenetic cell-type identification (opto-tagging) with tangential recordings in SC. (**a**) Schematic of light fiber position for optogenetic activation relative to the API. (**b**) The recording sites located in the upper visual layers of the SC (white dashed lines) are identified via visually evoked activity (left). Optogenetic activation can cover a large fraction of the SC tissue (right). The average visual and LED evoked activity along the recording sites is shown below the colormap. Gray vertical bars = light artifacts induced by the LED stimulation. VGAT-ChR2 mice were used in this example. (**c**) Receptive fields of recording sites with single units showing LED-evoked activity (blue). Recording sites of non-modulated single units are shown in gray. (**d**) Overlapping raster plot (black dots) and the corresponding activity (black line) during LED activation (blue) of a representative optogenetically modulated single neuron. (**e**) Individual recordings yield a large number of optogenetically activated SC cells.

We identified populations of inhibitory SC neurons by optotagging via Channelrhodopsin-2 (ChR2) activation in VGAT-ChR2 mice. Before the Neuropixels probe implantation (see above), an optic fiber (NA .66, 200 μm diameter, with a bare fiber tip, PlexBright, Plexon) connected to a blue LED module (PlexBright, Plexon) was implanted using a 20° angle tilted towards the AP axis (Fig. 5a). The fiber was inserted in the tissue prior to the Neuropixels probe implantation to decrease mechanical pressure and reduce drift of the Neuropixels probe caused by pressure through the fiber. The insertion was zeroed at the brain surface and the angled fiber was slowly lowered until reaching a depth of ~400μm. Following the fiber implantation, the Neuropixels probe was inserted perpendicular to the fiber as mentioned above along the AP axis in an ~18° angle relative to the horizontal plane. Short LED light pulses were given to evoke spiking activity in SC and to verify the proper alignment of the optic fiber and the Neuropixels probe. If no optogenetically evoked neuronal activity was observed, either the fiber was slightly lowered to a maximum depth of 700 μm or the electrode position was adjusted to ensure the alignment of the optic fiber with the probe. Once properly aligned, LED stimulation was applied to identify optogenetically-activated MUA and single units.

Using this combined approach, we could recruit optogenetically evoked neuronal activity across a large range of recording sites located within the SC (Fig. 5b). SC was identified via the visually evoked channels as described above (Fig. 5b, left). A precisely placed high-density probe inside SC together with a well aligned optic fiber can yield optogenetically evoked activity across a large fraction of the recording sites (up to 140 channels in the example shown in Fig. 5b), with RF locations of optogenetically modulated channels covering a large extent of the visual field (Fig. 5c). Since the Neuropixels probe is highly light-sensitive (Jun et al., 2017), the LED stimulation can cause light artefacts, transient voltage deflections, preferentially at the onset and offset of light pulses. Therefore, the LED onset (−2 to 6ms) and LED offset (−0.6 to 1 ms) were excluded from the data analysis (Bennett et al., 2019; Duan et al., 2021; Sans-Dublanc et al., 2021).

To estimate the number of single units that can be identified using this technique we performed spike sorting and manual curing using Kilosort2 (Pachitariu et al., 2016) and Phy (https://github.com/cortex-lab/phy). Our combined approach allows us to identify inhibitory cell type populations (45 identified cells in the example shown in Fig. 5) that consisted of ~38% out of the total recorded SC population, (Fig. 5d/e, 45 optogenetically modulated units out of 118 total SC units). In summary, combining the tangential recording configuration with orthogonally placed optical fibers can yield a high number of identified cells of a given type in SC.

## Discussion

Revealing how sensory information is represented and encoded on the population level within neural circuits of the brain is an important goal of neuroscience. While progress has been made to study populations of neurons in cortical circuits (de Vries et al., 2020; Froudarakis et al., 2014; Siegle et al., 2021; Stringer et al., 2019), less is known about how sensory stimuli are simultaneously processed in populations of neurons within subcortical circuits, e.g. the superior colliculus. This is partly due to the fact that subcortical structures are not easily accessible by imaging approaches which are the standard method of choice when it comes to population measures. Another limitation of the imaging technique is the field of view that only allows to study a subset of the retinotopy represented by the population. Here, we have introduced and described an approach that enables to study large populations of neurons in superior colliculus: tangential high-density electrode recordings

The high-density electrodes can capture the collective activity of high numbers of neurons. Commonly these electrodes are inserted in the brain from above, in a close to perpendicular angle. While vertical insertion angles capture neuronal activity e.g. across the different layers in cortex or superior colliculus simultaneously, the relatively thin structures restrict the measurement of large neural populations within single layers. Furthermore, the vertical insertion restricts the ability to record neuronal activity to a localized part of the visual field. How visual stimuli are simultaneously processed across the entire visual field cannot be studied using only vertical implantation angles and requires to record from a large extent of retinotopy. Our approach overcomes these limitations by recording along the horizontal plane of the superior colliculus, thereby simultaneously capturing the activity of neurons at various retinotopic locations. In addition, we demonstrate that the tangential recording approach can be combined with optogenetic tools to identify a high yield of neurons of a specific cell-type.

Recent studies have investigated the processing and encoding of visual information at the level of the retina and revealed important circuit mechanisms (Demb and Singer, 2015; Field and Chichilnisky, 2007; Pillow et al., 2008). However, how visual information is represented in neuronal populations within the SC, which receives direct retinal input, is less understood. Our approach allows to address these questions. Particularly, how are the different parts of the visual field, specifically the upper and lower visual field, simultaneously processed in populations of SC neurons? For example, center-surround interactions are ubiquitous in visual circuits, including SC. While center-surround interactions have been studied on the level of single neurons or small neural populations (Barchini et al., 2018), simultaneous measurements of neurons in center and surround have been challenging. Here, the tangential insertion overcomes this limitation by recording neurons simultaneously across a large region of visual space. This will enable to investigate center-surround interactions on the population level.

While we have presented here data from anesthetized mice, our approach is also applicable in awake head-fixed mice using the same procedure. This opens up opportunities to investigate how behaviorally relevant visual stimuli in the upper and lower visual field are simultaneously integrated within the SC circuitry and how the activity of the distinct circuits and neuronal populations of the SC relate to behavioral events. In particular, recent studies have indicated that SC plays a more prominent role in higher cognitive functions than previously thought (Basso et al., 2021). Combining the tangential recording approach with modern and sophisticated behavioral paradigms in mice has the potential to significantly advance our understanding on the role of SC during behavior.

A prominent feature of neural circuits within sensory systems is their organization in functional maps. The mouse SC was reported to be organized into a map of orientation (Ahmadlou and Heimel, 2015; Feinberg and Meister, 2015) and direction (Li et al., 2020), however, recent results challenge this conclusion (Chen et al., 2021). Tangential recordings have revealed important insights about the functional organization of the visual cortex, either by systematically advancing single electrodes (Hubel and Wiesel, 1977) or by simultaneously measuring neuronal activity using multi-electrode arrays (Kremkow et al., 2016). Here, we show that tangential Neuropixels recordings allow to characterize the functional responses in SC on a large scale and across a large extent of the SC circuitry. Thus, this method has the potential to answer and resolve the ongoing discrepancies of reports on the functional organization of mouse SC.

Finally, the tangential high-density approach described here is achievable, and thus is broadly applicable to study the processing of large populations in neural circuits across the brain. For example, future work could take advantage of a tangential insertion to target the horizontal dimension of the mouse cortex or the hippocampal formation which will reveal interesting new insights into population representations of the sensory environment in both allocentric and egocentric space.

## Conclusion

Tangential recordings with high-density electrode probes open up the possibility to study the population representation of sensory information across a large part of the visual field in awake behaving mice.

## Acknowledgement

We thank T. Lupashina for comments on the manuscript and T. Leva, and P. Schnepel for help with the Neuropixels recordings; J. Siegle and D. Denman for help with software (OpenEphys) and setup. We thank the whole Neuropixels community for their equipment and support and the Allen Institute for Brain Science for fostering high quality databases. This work was supported by the DFG Emmy-Noether grant KR 4062/4–1 (JK).

